# Structural and Functional Connectivity Predict the Effects of Direct Brain Stimulation on Memory

**DOI:** 10.64898/2026.03.17.712408

**Authors:** Qirui Zhang, Youssef Ezzyat, Ruoyi Cao, Sam S Javidi, Michael R Sperling, Michael J Kahana, Joseph I Tracy, Noa Herz

## Abstract

Intracranial stimulation can enhance episodic memory in humans; however, the behavioral effects vary substantially across individuals and stimulation sites. Here, we investigated whether the network embedding of a stimulation target, defined by MRI-based normative structural and functional connectivity, accounts for variability in stimulation-linked memory enhancement. We analyzed data from 50 adults with medically refractory epilepsy who underwent intracranial EEG monitoring and completed a verbal delayed free-recall task during stimulation of left temporal cortex sites across 61 sessions (39 closed-loop; 22 random). On average, closed-loop stimulation delivered during classifier-detected low-encoding states increased recall rates, whereas random stimulation produced no reliable benefit. Diffusion tractography from a normative database showed that sites yielding greater memory enhancement were characterized by stronger structural coupling to a distributed fronto-temporo-parietal network. Greater structure-function congruence with a normative verbal-encoding activation network predicted larger closed-loop memory benefit (Spearman ρ = 0.58, P < 0.0001). Functional connectivity exhibited overlapping trends but did not yield robust regional associations after permutation correction. Multivariate Partial Least Squares Structural Equation Modeling further identified stimulation mode, baseline memory, and a structural profile factor as independent predictors of memory enhancement, with no independent contribution of functional connectivity. These findings indicate that reliable stimulation-driven memory improvement depends not only on the timing of stimulation, but also on whether the stimulated target is structurally embedded within an encoding-relevant network scaffold.

**Significance statement:** Memory enhancement through direct brain stimulation holds substantial clinical promise, yet inconsistent outcomes have limited its therapeutic translation. This study shows that the effectiveness of closed-loop brain stimulation for memory improvement is determined by the structural network architecture of the stimulation target. Sites more deeply embedded within white-matter pathways connecting a distributed verbal encoding network yield the greatest mnemonic benefits when stimulation is delivered adaptively during poor encoding states. These findings establish a principled, network-based rationale for precision-guided neuromodulation: optimizing both the target’s structural embedding and the timing of stimulation delivery are necessary and complementary conditions for reliable, individualized memory enhancement.

## Introduction

Memory disorders constitute a major and growing public health challenge, affecting millions of individuals worldwide and contributing substantially to loss of independence and quality of life (1). Direct electrical stimulation of the human brain has emerged as a promising approach for modulating memory processes, as exemplified in patients with epilepsy who are implanted with intracranial electrodes as part of their clinical care. These indwelling electrodes provide a unique opportunity to investigate the neural mechanisms of memory and test causal interventions to restore cognitive function. While several studies have attempted to enhance memory through direct brain stimulation, the predominant outcome has been transient memory disruption rather than improvement, particularly when targeting the hippocampus or nearby mesial temporal lobe structures(2–5). More recent work suggests that targeting lateral temporal lobe regions can produce more reliable benefits, especially when stimulation is delivered in a state-dependent manner. Closed-loop approaches, in which stimulation is triggered by real-time neural signals predicting imminent memory failure, have provided some of the strongest evidence for memory rescue in humans (6, 7). Yet, even with closed-loop stimulation, substantial inter-individual variability persists, with some patients showing no benefit or even memory decrements.

Several factors have been proposed to account for this variability beyond stimulation timing, including anatomical target, stimulation parameters, and baseline memory ability(8–10). An increasingly compelling determinant, however, is the degree to which stimulation engages white-matter pathways that support communication across distributed memory circuits. White matter comprises axonal bundles that form the structural routes of inter-regional signaling(11). Because electrical stimulation can recruit axons near the stimulation site, stimulation delivered closer to major fiber pathways may propagate more effectively to distant nodes of a cognitive network.

Consistent with this mechanism, stimulation near mesial temporal lobe white-matter pathways has been reported to yield more favorable memory effects than stimulation confined to gray matter(11, 12), with similar gray-white differences reported for lateral temporal cortex (LTC) stimulation(13). Notably, recent closed-loop studies indicate that reliable memory enhancement is most evident when stimulated electrodes are proximal to major white-matter tracts, whereas superficial cortical stimulation farther from white matter shows weaker or absent benefits (14).

Beyond white-matter proximity, stimulation efficacy also appears to depend on functional interactions with memory-relevant circuits. Prior work has shown that stimulation targets yield greater memory enhancement when they are functionally connected to a patient’s memory network, as assessed using low-frequency intracranial EEG coherence (12). These findings raise the possibility that stimulation efficacy depends not on local anatomy, but on whether stimulation sites are embedded within the broader structural and functional architecture of the memory network. However, the relationship between functional connectivity and the underlying structural pathways that support it - and their joint contribution to stimulation outcomes - has not been systematically tested.

A network-level account of stimulation efficacy is further supported by evidence that episodic memory is implemented by distributed hippocampal–neocortical circuits, rather than isolated regions (15, 16). Stimulation has been shown to modulate both local and remote neural activity through engagement of large-scale networks, thereby shifting cognitive states(17–24).

Intracranial recordings further reveal consistent neural signatures of successful episodic memory encoding and retrieval, typically increases in high-frequency activity (>44 Hz) and decreases in alpha/beta power (8–30 Hz), distributed across LTC, medial temporal lobe, and prefrontal cortex(13, 25). Given the distributed nature of both episodic memory and stimulation-evoked effects, it is plausible that behavioral responses to stimulation are shaped by how strongly the stimulated region and its downstream pathways overlap with the circuits supporting an individual’s memory processes.

Despite growing recognition of network-level stimulation effects, memory-enhancing outcomes have not been consistently replicated, suggesting that a mechanistic account must move beyond stimulation timing and gross anatomical location alone. Given that proximity to white matter is an established predictor of stimulation efficacy, a critical open question is what mechanism links white-matter distance to behavioral outcome. Proximity per se is a geometric measure that does not specify which distributed circuits a stimulation site can actually engage; two sites equidistant from the gray-white boundary may differ substantially in their network architecture and, consequently, in the memory-relevant regions they can reach via fiber pathways. We therefore propose that it is the structural connectivity profile of the stimulation site, capturing which distributed regions are reachable through white-matter pathways and how strongly those regions are coupled to verbal encoding networks, that constitutes the operative mechanism linking white-matter embedding to behavioral outcome. Here, we directly test this hypothesis by examining whether network embeddings of stimulation sites account for variability in stimulation-related memory change. Specifically, we quantify the structural and functional connectivity of LTC stimulation sites and evaluate their alignment with a task-derived functional verbal-encoding network (Fig. 1). Using both random stimulation and closed-loop stimulation triggered during classifier-detected poor encoding states, we tested the hypothesis that memory benefits would be greatest when LTC targets are structurally coupled to, and aligned with, an encoding-relevant memory network. Finally, we jointly model the contributions of stimulation-site structural connectivity, functional connectivity, stimulation mode, proximity to major white-matter tracts, and baseline memory performance to delineate their independent and combined influences on stimulation efficacy.

**Figure 1.**
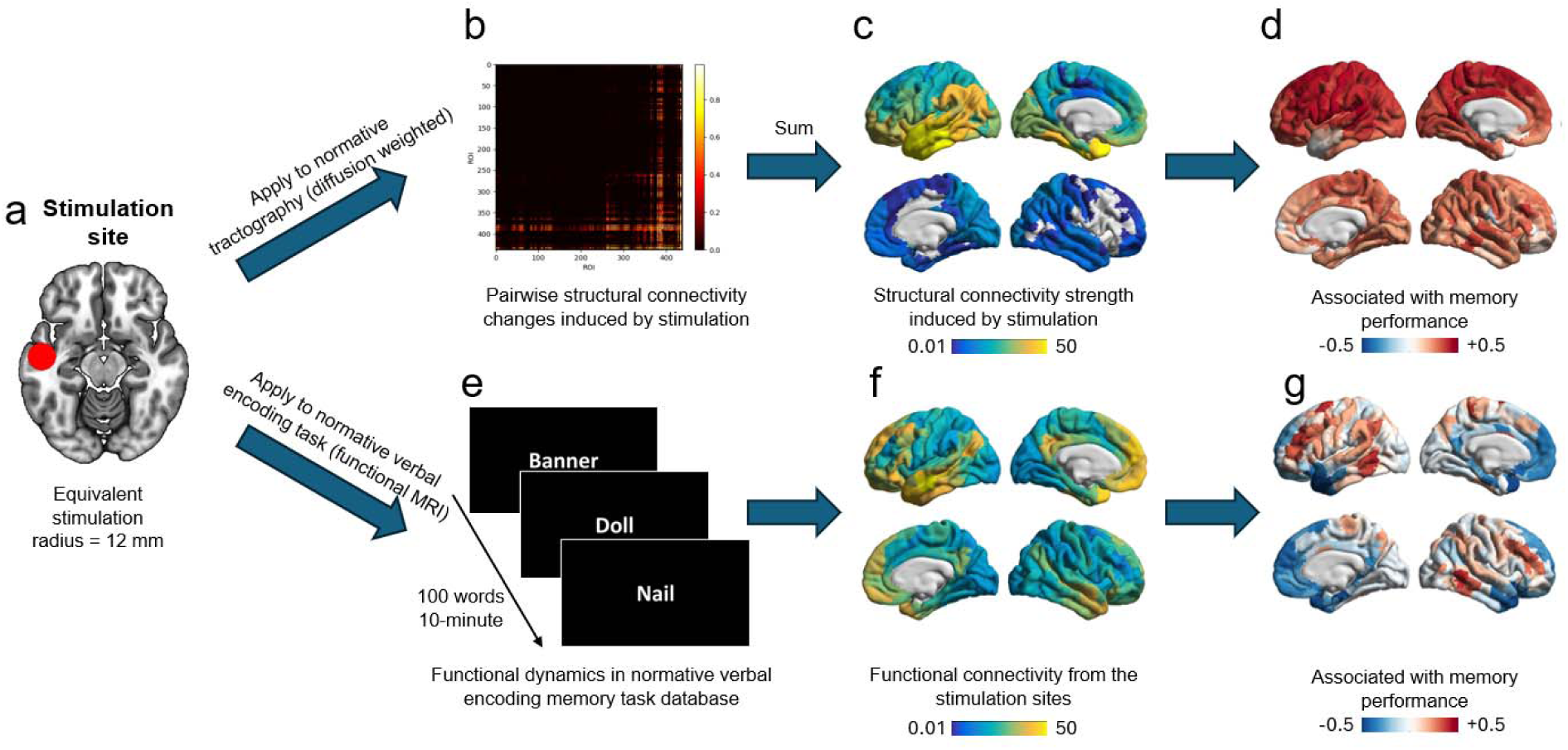
Experimental design and analytic framework. (a) Example of a stimulation site in the left lateral temporal cortex (LTC) from a representative participant, modeled as a 12 mm sphere representing the equivalent stimulation radius in MNI space. (b) Pairwise ChaCo (Change in Connectivity) ratio between the stimulation site and each brain region, reflecting the proportion of normative tractography streamlines connecting each region-pair that intersect the stimulation sphere. Warmer colors (yellow) indicate a higher proportion of intersecting streamlines, reflecting stronger structural coupling; cooler colors (red/black) indicate weaker or absent structural coupling. . (c) Overall structural connectivity strength in the same participant, obtained by summing pairwise connectivity changes across all target brain regions, shows the distributed structural pathways engaged by stimulation. (d) Depiction of the stimulation-induced structural connectivity profiles associated with memory performance computed across all participants. (e) Schematics of verbal encoding fMRI paradigm used to capture memory-related functional dynamics. (f) Depiction of functional connectivity from the same individualized stimulation site during the verbal-encoding fMRI task, illustrating the distributed functional pathways engaged by stimulation. (g) Depiction of stimulation-induced functional connectivity profiles associated with memory performance across all participants.

## Results

### The Lateral Temporal Cortex is a Reliable Target for State-Dependent Memory Enhancement

Based upon previous work (6, 7, 14, 26, 27) showing that the stimulation of left LTC was associated with reliable verbal memory enhancement, we focused our analyses on the left LTC to characterize the structural connectivity pathways associated with successful memory modulation via electrical stimulation. Importantly, previous studies have shown that memory enhancement at this site critically depends on the stimulation methodology - specifically, whether stimulation is delivered in a closed-loop or random fashion. Accordingly, we preserved this distinction in all subsequent structural pathway and behavioral analyses.

In this paradigm, stimulation was delivered during a free recall task designed to detect periods of poor memory encoding, operationalized as neural states predictive of subsequent recall failure.

Two stimulation approaches were employed: (i) a closed-loop condition, in which stimulation was triggered in real time based on these neural biomarkers, and (ii) a random condition, in which stimulation was delivered at randomly selected events independent of encoding state (see Fig. 2a).

**Figure 2.**
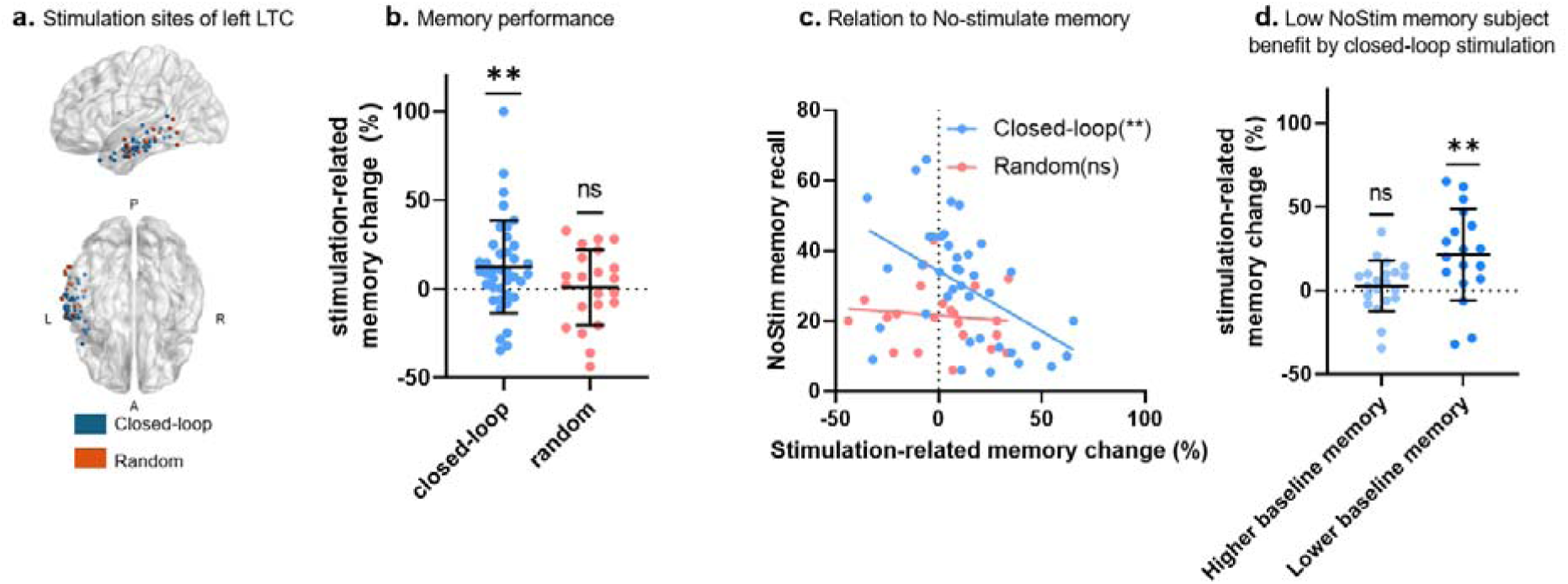
Closed-loop stimulation improves memory performance. (a) Anatomical distribution of stimulation sites within the left LTC for closed-loop (blue) and random (red) conditions across participants. (b) Stimulation-related memory change (Δ recall, %) for closed-loop (blue) and random (red) conditions. Each dot represents an individual participant’s memory change score (Δ). Horizontal lines denote the mean, and error bars indicate the standard deviation. Closed-loop LTC stimulation significantly improved recall performance relative to baseline memory recall, whereas random stimulation had no effect. (c) Scatter plot of stimulation-related memory change (Δ recall, %) as a function of baseline recall performance, shown separately for closed-loop (blue dots) and random (red dots) participants. In the closed-loop condition, stimulation-related memory improvement was inversely correlated with baseline memory recall performance. (d)Stimulation-related memory change (Δ recall, %) in closed-loop participants stratified by baseline memory performance into low- (dark blue) and high-baseline (light blue) subgroups. Closed-loop stimulation selectively benefited subjects with lower baseline memory, with no reliable effect in those with higher baseline memory; ns, not significant (P > 0.05); * P < 0.05; ** P < 0.01.

Memory enhancement was quantified as the difference in recall performance between lists containing stimulation and baseline (non-stimulated) lists. Closed-loop stimulation of the left LTC produced a significant improvement in recall performance relative to baseline (Δ = 12.4% ± 4.18 SEM; *t*(38) = 2.96, *P* = 0.005; Fig. 2b), demonstrating that targeted stimulation during suboptimal encoding states enhances subsequent memory retrieval. In contrast, random stimulation produced no significant change in recall performance (Δ = 0.92% ± 4.52 SEM; *t*(21) = −0.20, *P* = 0.83). These findings replicate and extend prior demonstrations of memory enhancement via closed-loop LTC stimulation (6, 7, 14, 26, 27).

We next examined whether stimulation-related memory change depended on an individual’s baseline memory performance. In the closed-loop condition, stimulation-induced memory improvement was significantly and negatively correlated with baseline recall performance (r = −0.48, df = 37, P = 0.0017), indicating that individuals with poorer baseline memory derived greater benefit from stimulation. No such relationship was observed in the random stimulation group (r = −0.10, df = 20, P = 0.63; Fig. 2c). To further characterize this effect, closed-loop participants were stratified into low- and high-baseline memory subgroups using a threshold corresponding to mean baseline performance (threshold = 30%, corresponding to the mean performance). Closed-loop stimulation selectively enhanced recall in participants with low baseline memory performance (Δ = 21.54% ± 6.44 SEM; *t*(20) = 3.34, *P* = 0.004), while producing no reliable effect in participants with higher baseline memory (Δ = 2.80% ± 3.30 SEM; *t*(17) = 0.84, *P* = 0.40)). A two-sample t-test also confirmed that subjects with poor baseline memory exhibited significantly greater stimulation-related improvement than those with higher baseline memory (*t*(37) = 2.69, *P* = 0.010, Fig. 2d).

### Structural Connectivity Defines Memory-Enhancing Stimulation Sites

Building on prior evidence that stimulation proximity to white matter pathways predicts memory outcomes(14), we extended our analyses to diffusion tractography to test whether the structural connectivity profile of stimulated temporal lobe sites could account for the individual differences observed in stimulation-related memory outcomes. Notably, as shown in Fig. 2b, even under closed-loop stimulation, a subset of participants failed to exhibit memory enhancement and, in some cases, demonstrated memory impairment, highlighting the need to identify circuit-level determinants of stimulation efficacy.

For each stimulation site, we first constructed a normative structural connectivity profile using diffusion tractography from healthy participants (HPs, see Methods). We then quantified the relationship between each site’s structural connectivity profile and stimulation-induced memory change. In the closed-loop stimulation condition (df = 37), sites associated with greater memory improvement exhibited markedly stronger connectivity to a distributed network encompassing frontal, parietal, superior temporal, and mid-cingulate regions (Fig. 3a). No such relationships were observed in the random stimulation condition (df = 20, Fig. 3b). These findings indicate that for close-loop stimulation , memory enhancement depends on whether the stimulation site is structurally embedded within a specific distributed cortical network. The spatial organization of this network closely corresponds to a canonical verbal encoding system, comprising regions known to support episodic memory through complementary cognitive operations. These include the left LTC, which supports higher-order semantic representations and lexical access (28, 29); the left inferior frontal gyrus, which mediates controlled semantic retrieval, selection, and articulatory–phonological processes during encoding (16, 30); the left dorsolateral prefrontal cortex, which contributes strategic working-memory operations and top-down attentional control (31); and posterior parietal cortex, which supports the integration of contextual and mnemonic information and facilitates episodic retrieval processes (32).

**Figure 3.**
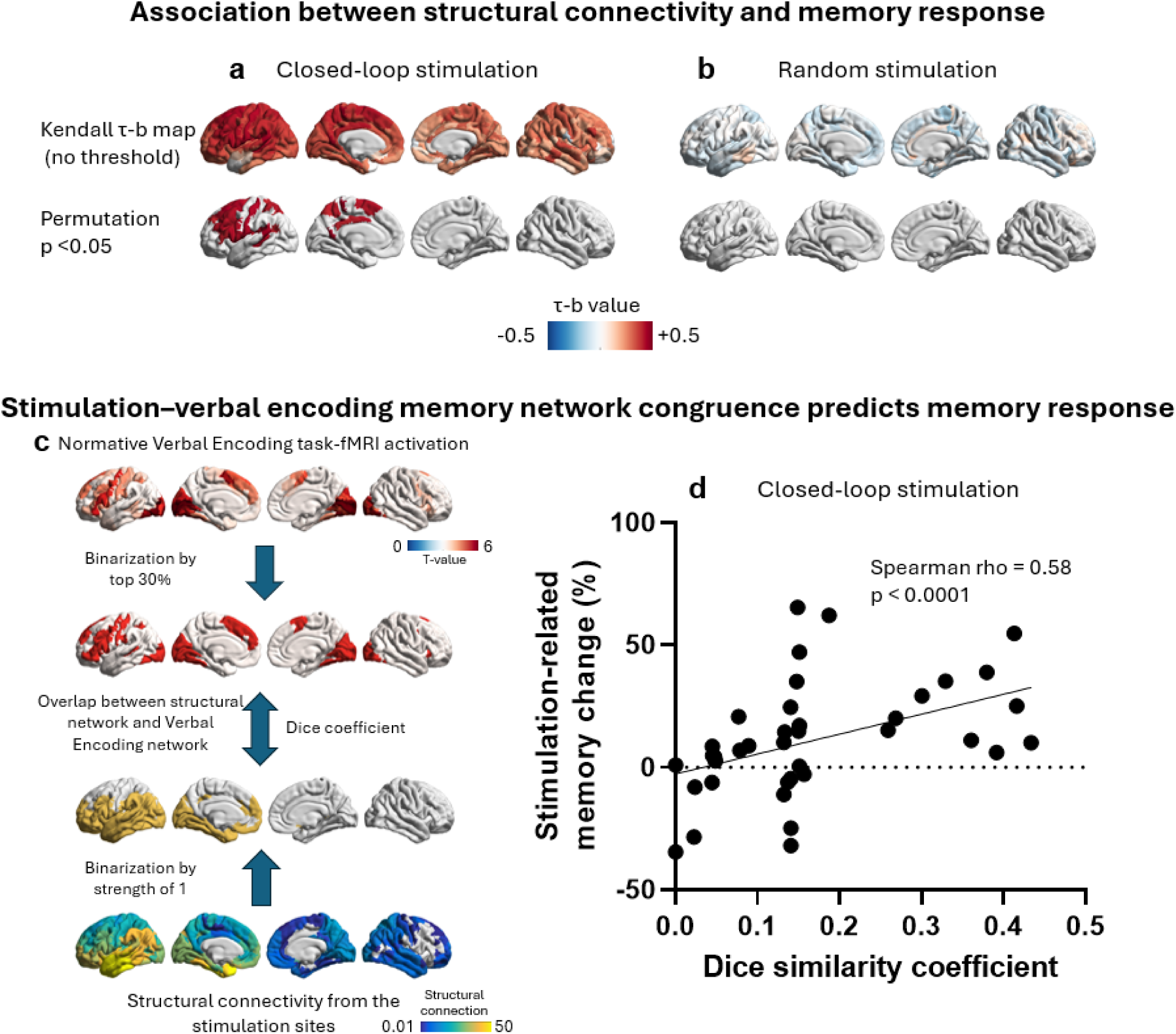
**Memory-enhancing are defined by their structural connectivity profile**. (a) Brain surface maps display the Kendall τ-b correlation coefficient computed, for each of 438 cortical parcels, between that parcel’s normative structural connectivity to the stimulation site and stimulation-related memory change across participants. Positive correlations (warm colors) indicate that stronger structural coupling between a given parcel and the stimulation site was associated with greater memory improvement. The bottom panel displays only parcels surviving permutation-based family-wise error correction (*p* < 0.05, 5,000 iterations); the top panel shows the full uncorrected map for comparison. In the closed-loop condition (df = 37), stimulation sites associated with greater memory improvement exhibited stronger structural connectivity to distributed frontal, parietal, superior temporal, and mid-cingulate regions. No subcortical or cerebellar regions showed significant associations and are therefore not displayed. (b) In the random stimulation condition (df = 20), no reliable relationship was observed between structural connectivity profiles and memory performance. (c) Illustration of the structure-function congruence analysis. A normative verbal encoding network was derived from task-based functional MRI data collected in HPs. For each stimulation site, a participant-specific structural connectivity map was generated using diffusion tractography. Both maps were thresholded (top 30% of the functional verbal encoding network and strong structural connectivity values > 1) and binarized prior to computing their spatial overlap. The Dice similarity coefficient quantified structure-function congruence for each site. (d) Scatter plot showing the relationship between structure–function congruence (Dice similarity coefficient; x-axis) and stimulation-related memory change (%; y-axis), where each dot represents a single subject . Higher structure– function congruence was strongly associated with greater stimulation-related memory improvement in the closed-loop condition (*p* < 0.001). This association remained robust across a wide range of functional-network and structural-connectivity thresholds. In contrast, no such relationship was observed in the random stimulation group (Supplementary Fig. S1).

Together, these results suggest that stimulation is most effective when its structural projections engage a broader functional system supporting verbal memory encoding. To directly test this structure-function coupling, we derived a normative verbal encoding network from task-based functional MRI data acquired in a local cohort of HPs performing a word-list encoding task comparable to the stimulation paradigm. Across a broad range of verbal encoding network and structural-connectivity thresholds, a robust positive association between structure–function congruence and stimulation-related memory improvement was consistently observed in the closed-loop stimulation condition (Supplementary Fig. S1). To illustrate a representative operating point, Fig. 3c,d highlights results obtained using thresholds retaining the top 30% of voxels within the verbal encoding network and structural connectivity strength greater than 1. At this representative threshold, greater structure-function congruence was strongly associated with larger memory enhancement following closed-loop stimulation (Spearman ρ = 0.58, df = 37, P < 0.0001). These findings indicate that stimulation targeting core regions of the memory network that are embedded within strong structural connectivity yields the strongest association with memory enhancement. In contrast, no significant relationship was observed in the random stimulation group at any of the thresholds tested (Supplementary Fig. S1). These findings provided compelling evidence that when the stimulation signal (i.e., closed-loop) is propagated by structural connections embedded within a verbal encoding network, the memory enhancement effect is stronger. Conversely, stimulation delivered outside this network - or delivered independently of encoding state - fails to reliably improve memory performance, underscoring the importance of both network embedding and state-dependent targeting for effective neuromodulation.

### Functional Connectivity Profiles Associated with Memory-Enhancement

Having identified structural network embedding as robust predictors of closed-loop memory benefit (Fig. 3), we next asked whether functional coupling alone provides a complementary predictor of stimulation efficacy. Specifically, using task-based fMRI data from HPs, we derived normative whole-brain functional connectivity profiles for each stimulation site (see Methods) and tested whether stronger site-to-network functional connectivity was associated with stimulation-related memory change.

Across both closed-loop (df = 37) and random stimulation (df = 20) conditions, functional connectivity showed positive associations with memory change in regions that broadly overlapped those identified by structural connectivity analyses (e.g., bilateral dorsolateral prefrontal cortex, lateral temporal cortex, and angular gyrus; Fig. 4). However, none of these regional stimulation-site associations survived permutation-based multiple-comparison correction. At an uncorrected threshold (p < 0.05), the closed-loop condition exhibited a distributed pattern of positive correlations consistent with the canonical verbal encoding network (Fig. 4a), whereas no such effects were observed in the random stimulation condition even before correction (Fig. 4b). Together, these findings indicate that although functional connectivity patterns qualitatively mirrored the structural connectivity results, functional coupling alone did not provide a statistically robust predictor of stimulation-induced memory enhancement.

**Figure 4.**
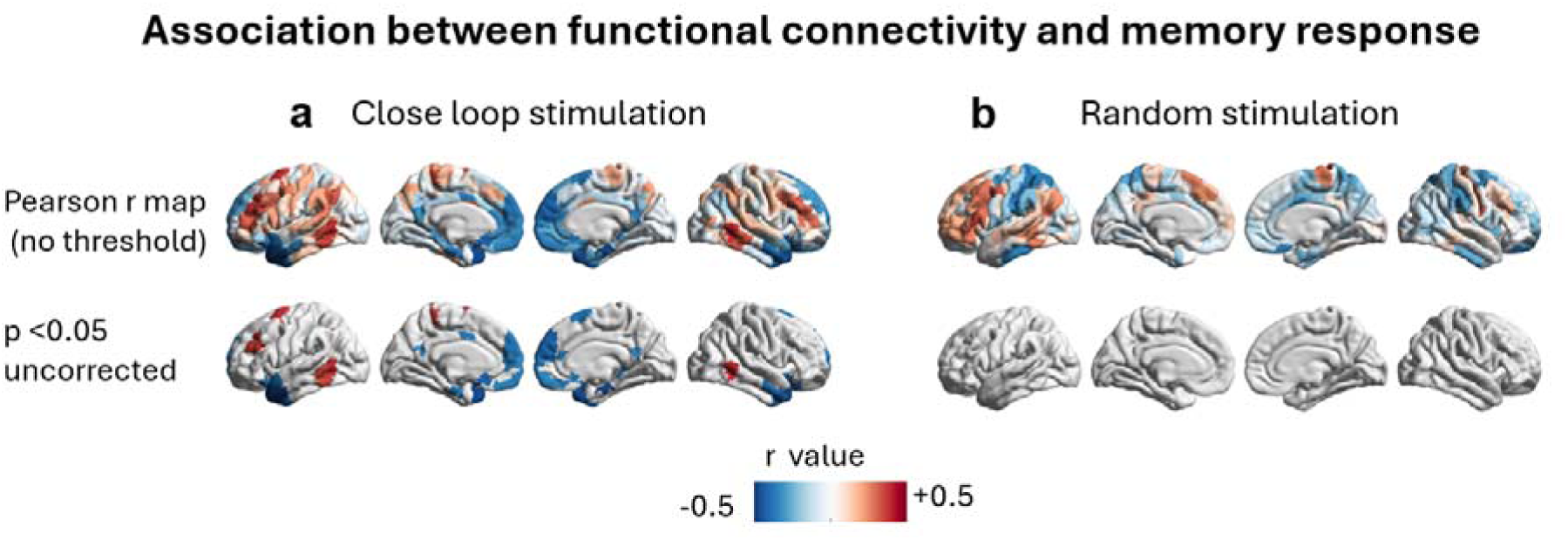
Functional connectivity profiles associated with stimulation-related memory change. In both panels, color indicates the Pearson correlation coefficient between each region’s normative functional connectivity to the stimulation site and stimulation-induced memory change across participants (a) In the closed-loop condition (df = 37), stimulation-related memory improvement showed positive (uncorrected) associations with functional connectivity for the dorsolateral prefrontal cortex, LTC, and angular gyrus regions (p < 0.05, uncorrected). No subcortical or cerebellar regions met this threshold and, therefore, are not displayed. None of the above functional connectivity-memory change associations survived permutation-based multiple-comparison correction. (b) In the random stimulation condition (df = 20), no regions displayed statistically reliable functional connectivity/memory change associations, even at an uncorrected threshold (p < 0.05).

### Modeling Multiple Determinants on Stimulation-Induced Memory Enhancement

These findings indicate that stimulation-induced memory enhancement is influenced by multiple factors, including the structural network embedding of the stimulation site, baseline memory performance, and the mode of stimulation. In the following analyses, we aimed to disentangle the unique and shared contributions of structural connectivity relative to these other determinants, and to test whether its effects interact with stimulation mode and individual baseline memory capacity.

To evaluate these relationships, we applied partial least squares structural equation modeling (PLS-SEM). This framework enables dimensionality reduction followed by regression on latent constructs to model and quantify relationships between multiple independent variables. Predictor variables included five structural and five functional connectivity measures derived from the five brain regions most strongly associated with stimulation-related memory change in the closed-loop condition, baseline memory performance, stimulation mode (closed-loop versus random), the WMP measures, and stimulation-related memory change as the outcome variable. Path coefficients were estimated using 5,000 bootstrapped samples. The structural and functional connectivity, and WMP measures comprised separate sets of indicators, with each decomposed into one latent factor. We first modeled the separate effects of the structural and functional connectivity latent factors, along with stimulation mode as a moderating factor, on memory enhancement (see PLS-SEM models depicted in Fig. 5a). We then modeled the separate effects of the structural and functional connectivity latent factors, along with the WMP latent factor, on memory enhancement (see PLS-SEM models depicted in Fig. 5b). Lastly, we constructed a multivariate model including the structural and functional connectivity latent factors, along with stimulation mode and baseline memory performance, for the prediction of memory enhancement (PLS Multivariate SEM regression model).

**Figure 5.**
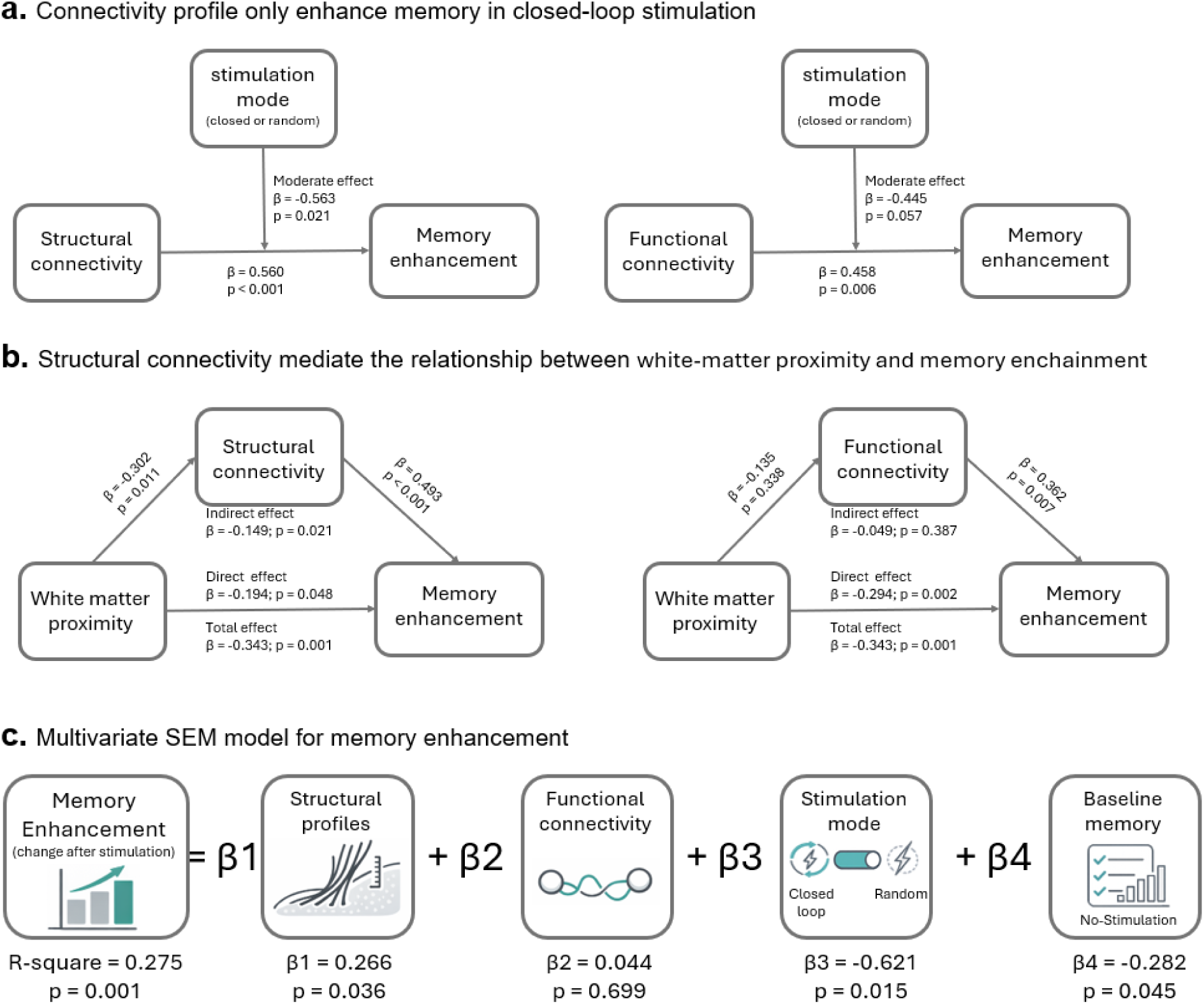
Partial least squares structural equation models. (a) Connectivity predicts stimulation-related memory enhancement in the closed-loop, but not the random, stimulation condition. Left, structural connectivity model; right, functional connectivity model. Standardized path coefficients (β) and p values are shown for both direct and moderating effects. (b) Structural connectivity mediates the effect of WMP on memory enhancement, whereas functional connectivity does not. Left, structural connectivity mediation model; right, functional connectivity mediation model. Standardized path coefficients (β) and p values are shown for both direct and indirect (mediated) effects. (c) Independent and joint contributions of multiple predictors to stimulation-related memory outcomes. Shown are standardized path coefficients (β) and p values from a multivariate PLS-SEM model quantifying the unique and combined influences of stimulation mode, baseline memory performance, structural profile measures, and functional connectivity on memory enhancement.

Structural Connectivity Predicts Memory Enhancement Specifically During Closed-Loop Stimulation: Structural connectivity robustly predicted stimulation-related memory enhancement, with effects that were specific to closed-loop stimulation. Structural connectivity significantly predicted stimulation-related memory improvement (β = 0.560, p < 0.001), and this relationship was significantly moderated by stimulation mode (β = −0.563, p = 0.021). This moderation indicates that the relationship between structural connectivity and memory enhancement depended on stimulation mode. Specifically, the positive association between structural connectivity and memory enhancement was present only in the closed-loop stimulation condition. Functional connectivity also predicted memory improvement (β = 0.458, *p* = 0.006), demonstrating only a statistical trend for the potential moderating effect of stimulation mode (β = −0.445, *p* = 0.057). This pattern is consistent with the results reported above, indicating that connectivity-memory relationships were strongest and most reliable in the closed-loop condition (Fig. 5a).

Structural Connectivity Accounts for the Behavioral Advantage of White-Matter Proximity: Structural connectivity, but not functional connectivity, partially accounted for the behavioral advantage conferred by closer proximity of stimulation to white matter. Shorter distance to the gray-white matter boundary was significantly associated with stimulation-related memory improvement (total effect: β = −0.343, *p* = 0.001). To determine whether this relationship operated through structural connectivity, we evaluated a mediation model. In this model, WMP significantly predicted structural connectivity (β = −0.302, p = 0.011), and structural connectivity strongly predicted memory enhancement (β = 0.493, p < 0.001). When structural connectivity was included in the model, the direct effect of WMP on memory improvement was attenuated but remained significant (β = −0.194, p = 0.048). The resulting indirect effect from WMP to memory enhancement via structural connectivity was significant (β = −0.149, p = 0.021), indicating that approximately 43% of the total effect of WMP on memory enhancement was transmitted through structural connectivity. Model comparison further supported the inclusion of structural connectivity. The extended model demonstrated improved fit according to Bayesian Information Criterion (Δ = 7.56), representing strong evidence in favor of the model including structural connectivity (33, 34). These results show that affiliation with the memory network increase stimulation efficacy on memory beyond white matter proximity alone. Together, these findings indicate that structural network embedding is a key mechanism linking stimulation geometry to behavioral outcomes. In contrast, functional connectivity did not mediate the relationship between WMP and memory enhancement (β = −0.049, p = 0.387; Fig. 5b), underscoring the specificity of white-matter-dependent effects to structural, rather than functional, network architecture.

Multivariate Model of Stimulation-Related Memory Outcomes: To evaluate the independent and joint contributions of multiple measured factors to stimulation-related memory outcomes, we implemented a multivariate PLS-SEM approach. Predictors included a structural profile latent factor, derived from the joint decomposition of structural connectivity and WMP due to their strong covariance; a functional connectivity latent factor; stimulation mode (closed-loop vs. random); and baseline memory performance (baseline memory recall scores). The model explained 27.5% of the variance in stimulation-related memory change (R² = 0.275, p = 0.001). Stimulation mode emerged as the strongest predictor, consistent with prior results (β = −0.621, p = 0.015), followed by baseline memory performance (β = −0.282, p = 0.045). The structural profile factor also independently predicted memory enhancement (β = 0.266, *p* = 0.036), confirming that structural network embedding contributes unique variance beyond stimulation mode and baseline memory ability. In contrast, functional connectivity did not independently predict memory outcomes (β = 0.044, p = 0.699) (Fig. 5c).

## Discussion

Despite growing interest in direct brain stimulation as a therapeutic tool for memory impairment, its effects remain highly variable across individuals. Here, we show that successful memory enhancement via electrical stimulation is governed by network embedding and brain state, rather than stimulation timing or target location alone. Specifically, we demonstrate that closed-loop stimulation delivered during poor encoding states yields robust memory improvement when applied at sites proximal to white matter tracts that are structurally connected to the verbal memory network. This suggests that memory enhancement is maximized when stimulation can propagate along anatomical pathways into distributed memory circuits. These results establish network embedding and state-dependent delivery as core principles for precision-guided neuromodulation of human memory.

The present findings replicate and extend prior work demonstrating the efficacy of closed-loop stimulation for memory enhancement (6, 7, 12). We demonstrate that closed-loop stimulation of left LTC significantly improves recall when applied during poor encoding states, whereas random (open-loop) stimulation produces no reliable benefit. This timing likely allows stimulation to interrupt an inefficient encoding state and re-engage neural dynamics associated with successful memory formation.Closed-loop stimulation may act by reinstating or amplifying these network-wide signatures via structural connections, thereby “rescuing” functional network states and promoting memory storage. Consistent with this interpretation, recent work demonstrates that closed-loop LTC stimulation phase-synchronized to hippocampal theta produces persistent increases in cortico-hippocampal network connectivity, an effect absent when stimulation is delivered without phase-locking (5), further underscoring that temporal precision—whether defined by encoding state or oscillatory phase—is critical for effective memory network engagement.

Beyond stimulation timing, our results indicate that the structural connectivity profile of a stimulation site is a principal determinant of whether, and to what extent, electrical stimulation enhances memory. In the closed-loop condition, stimulation sites associated with larger behavioral gains exhibited markedly stronger structural coupling to a distributed set of frontal, parietal, superior temporal, and mid-cingulate regions, whereas the same analyses yielded no reliable effects in the random-stimulation condition. Mechanistically, these findings are consistent with the idea that stimulation engages axons near the stimulation site and thus can influence remote nodes of a cognitive network most effectively when the target is coupled to the relevant white-matter architecture(17, 19).

Critically, the relationship between structural connectivity and memory improvement was not merely anatomical “reach,” but structure-function alignment: stimulation sites whose structural projections overlapped more strongly with a normative verbal-encoding activation network produced the greatest closed-loop benefit. The robustness of this effect across a wide range of structural and functional threshold choices argues against an artifact of arbitrary parameterization and instead supports a genuine coupling principle, namely, that stimulation is most effective when its anatomical propagation routes converge on regions actively engaged in the targeted cognitive operation (6, 14). This pattern suggests that stimulation benefits are not simply a function of local anatomy, but instead depend on whether the target is structurally embedded within a larger memory-relevant circuit capable of supporting long-range signal propagation (14, 19, 24). Such a network-embedding account is further strengthened by the observation that the cortical distribution of structurally predictive regions closely resembles a canonical verbal encoding system, implicating complementary operations spanning semantic representation and lexical access, controlled retrieval and selection, executive control, and integrative mnemonic processing(16, 28–32). Within this framework, structural connectivity provides the causal substrate for stimulation-driven perturbations to spread, while the functional encoding network defines the computational context in which those perturbations can translate into improved behavior(19, 24). This interpretation also aligns with our WMP finding, stimulation sites closer to the gray-white boundary yielded greater memory benefit, an effect partially mediated by structural connectivity. Together, these results support the notion that stimulation is most efficacious when it can access tract pathways linking distributed encoding circuitry, rather than acting locally within gray matter alone (14, 17).

The present findings extend and mechanistically elaborate prior evidence that white matter proximity is a key anatomical determinant of stimulation efficacy. Previous work established that stimulation delivered closer to major fiber tracts yields greater memory benefit than stimulation confined to gray matter (14), yet the mechanism linking physical proximity to behavioral outcome remained unspecified. Our mediation analysis provides a causal account of this relationship: white matter proximity influences memory enhancement not as a direct physical determinant, but by indexing the degree to which a stimulation site is embedded within structurally relevant connectivity architecture. Specifically, structural connectivity accounted for approximately 43% of the total effect of white matter proximity on memory improvement, while retaining independent predictive value beyond proximity alone. This dissociation is theoretically important: two sites equidistant from the gray–white matter boundary can differ substantially in their structural connectivity profiles and, correspondingly, in their capacity to drive distributed network engagement. Proximity to white matter is thus best understood as a heuristic marker for network access, rather than the operative mechanism itself. The identification of structural connectivity as the mediating factor shifts the conceptual framework from geometric to topological: it is not how close a stimulation site is to white matter, but which white matter pathways it can recruit, and whether those pathways link the site to encoding-relevant circuitry, that ultimately governs stimulation efficacy.

Although functional connectivity showed broadly aligned qualitative patterns with the structural results, it did not independently predict memory outcomes. A plausible explanation is that fMRI-based functional connectivity emphasizes state-dependent coupling and is therefore more vulnerable to moment-to-moment variability, measurement noise, and person-specific reorganization of memory networks in epilepsy (35–37). By contrast, structural connectivity provides a more stable constraint on whether stimulation can consistently recruit memory-relevant regions, which may be especially important when stimulation is temporally gated to periods of heightened encoding vulnerability, as in closed-loop paradigms. Our functional-connectivity results also reflect a different operationalization than prior intracranial EEG work, where functional connectivity was quantified as low-frequency coherence between contacts within individualized memory networks defined from each patient’s free-recall performance (14). Here, functional connectivity was estimated from fMRI in healthy controls during a memory task, which may be less sensitive to low-frequency oscillatory dynamics and to individual-specific network architecture. Because normative network definitions can be particularly limiting in epilepsy, where functional organization may be substantially reorganized, individualized functional mapping may be required for functional connectivity to achieve comparable predictive power(38, 39). Together, these considerations support the conclusion that when normative memory networks are used, structural connectivity provides a more reliable and generalizable predictor for stimulation target selection in epilepsy patients.

The present findings also explain why random (i.e., open-loop) stimulation fails to reliably exploit individual memory network structure. Random stimulation is delivered without regard to the brain’s instantaneous encoding state and therefore often occurs when memory networks are already operating efficiently or are transiently disengaged from encoding-related processing.

Under these conditions, even stimulation applied to structurally well-connected sites may fail to interact constructively with the memory network. Moreover, without state-dependent gating, stimulation may be injected into network configurations that are functionally irrelevant to memory encoding, reducing the likelihood that structural pathways involved in memory are effectively recruited (6, 7). Thus, random stimulation lacks the temporal and spatial precision required to harness individual network architecture, rendering its effects weak or inconsistent despite comparable anatomical placement.

From a translational standpoint, our results strongly motivate personalized, network-guided neuromodulation strategies(8–10). Our multivariate modeling demonstrates that stimulation mode (closed-loop versus open-loop), baseline memory ability, and structural network properties independently predict memory enhancement. Future interventions must therefore optimize both when stimulation is delivered and where it is applied. Closed-loop systems can be used to align stimulation with optimal temporal windows defined by neural state(6, 7), while individualized structural connectomes can guide target selection toward regions deeply embedded within memory circuits and proximal to critical white-matter pathways(14, 19, 40). In practice, this implies that neuroimaging modalities such as diffusion MRI, combined with electrophysiological profiling, could be used to map individual memory networks prior to intervention and inform patient-specific stimulation strategies.

One important limitation of the present study is that structural and functional connectivity profiles were derived from high-quality normative datasets rather than from individualized patient-specific connectomes(40–43). While this approach enables robust embedding of stimulation sites within a canonical memory network framework, it does not capture individual variability in network reorganization, which is particularly relevant to epilepsy. Chronic seizure activity can profoundly alter connectivity patterns, leading to atypical or redistributed memory networks (44). In such cases, stimulation of a site traditionally considered a memory hub, but disconnected from the reorganized network, may be ineffective or even detrimental. Conversely, targeting regions that retain or acquire compensatory connectivity to memory circuits may yield greater benefit. The fact that our functional (fMRI) memory network measures, which were based on normative memory connectivity, did not independently predict memory enhancement suggests that, indeed, person-specific, atypically organized memory networks may be at work in our data. This raises a question about whether person-specific memory networks bear a reliable relationship to enhancement. Even further, it raises the possibility that capturing the alignment of person-specific structural and functional memory networks may foster strong enhancement effects. Understanding how individual-specific network reorganization shapes stimulation outcomes remains a critical challenge. As computational frameworks such as virtual brain twins continue to evolve, future work may be able to model stimulation-induced perturbations of whole-brain dynamics at the individual level (45–48), enabling more precise and mechanistically grounded neuromodulation strategies that would also allow assessment the role played by underlying structural connectivity.

In summary, this study identifies structural network embedding as a central determinant of stimulation-related memory enhancement and demonstrates the superiority of closed-loop stimulation for engaging memory networks in a state-dependent manner. By integrating network neuroscience with adaptive stimulation paradigms, our findings provide a principled framework for translating cognitive network theory into effective, individualized clinical interventions, pointing toward a future of precision-guided memory enhancement.

## Materials and Methods

### Participants

A total of 50 adults (mean age 38.3 ± 11.95 years; range 19-64 years; 28 male) with medically refractory epilepsy were enrolled. All participants underwent intracranial electroencephalographic monitoring as part of their presurgical evaluation for seizure localization (6, 7, 14, 26, 27). Electrode implantation and the selection of stimulation targets were determined solely by the treating clinical teams based on each patient’s seizure focus and surgical planning needs.

Stimulation was delivered to sites within the left LTC during memory testing (Figure 2A). Across the cohort, 61 unique stimulation sites were examined: 41 patients were stimulated at a single site, seven patients at two distinct sites, and two patients at three distinct sites. Only one site was stimulated during any given experimental session. During stimulation, participants performed a word list delayed free recall task with either semantically categorized or semantically unrelated words. In total, 61 stimulation sessions were conducted, comprising 39 closed-loop (35 participants; mean age 41.4 ± 12.15 years; range 19-64 years; 20 male) and 22 random (15 participants; mean age 32.75 ± 9.19 years; range 20-48 years; 8 male) stimulation trials.

Data were collected as part of a coordinated, multicenter research program investigating the effects of direct electrical stimulation on human memory processes. Clinical data collection was carried out at the University of Texas Southwestern Medical Center (Dallas, TX), Dartmouth– Hitchcock Medical Center (Lebanon, NH), Thomas Jefferson University Hospital (Philadelphia, PA), Emory University Hospital (Atlanta, GA), Mayo Clinic (Rochester, MN), the Hospital of the University of Pennsylvania (Philadelphia, PA), and Columbia University Medical Center (New York, NY). 40 of the current participants were included in a previous publication(14), and included only subjects who received stimulation to the left LTC. The remaining 10 patients were included on another publication(6) and included patients receiving random stimulation to the left LTC; however, all analyses and results reported here are novel. Supplementary table 1 shows patients’ numbers and demographic characteristics. The study was approved by the Institutional Review Board at each participating center, and written informed consent was obtained from all participants.

### Stimulation and memory performance

**Verbal memory task:** Participants were instructed to study a list of 12 words, followed by a short distraction period (20 seconds of a math distractor task in the form of A+B+C=?, where A,B,C are randomly chosen integers ranging from 1-9). Participants were then given 30 seconds to recall as many items from the preceding list, in any order. Vocal responses were digitally recorded and later manually annotated for analysis. Participants completed one of two variations of the verbal free recall task. In the standard free recall task, words were selected randomly from a pool of common nouns (https://memory.psych.upenn.edu/Word_ Pools). In the categorized free recall task, the word pool was constructed from 25 semantic categories (e.g., fruit, insects, and vehicles). Words were randomly drawn from three semantic categories per list and presented in same category pairs (e.g., “Ant”, “Bee”, “Apple”, “Pear”, etc.,). Each session consisted of 25 lists of this encoding-distractor-recall procedure.

**Stimulation Procedure:** Prior to each stimulation session, the safe amplitude for stimulation was determined while a neurologist monitored for after discharges. Stimulation amplitude started at 0.5 mA and was increased in steps of 0.5 mA, up to a maximum of 1.5 mA for depth contacts and 3.5 mA for cortical surface contacts. These maximum amplitudes were chosen to be below accepted safety limits for charge density (Shannon 1992). Stimulation was delivered in a bipolar fashion, between pairs of adjacent electrode contacts. Stimulation used charge-balanced biphasic rectangular pulses (pulse width = 300 μs) for a duration of 500 ms. Stimulation frequency varied between 50, 100, or 200 Hz, with a single frequency value per subject.

<u>Closed-loop stimulation:</u> The closed-loop cohort consisted of individualized memory classifiers to control the timing of stimulation in response to brain activity. At least three record-only sessions were collected from an individual patient to train a classifier to discriminate between words that were recalled and those that were not, using an L2-penalized logistic regression model. The model used spectral power features computed from intracranial recordings during the encoding window (0–1,366 ms after word onset), summarized across a small set of logarithmically spaced frequencies spanning approximately 3–180 Hz (Morlet wavelets, wave number = 5). For a subset of participants, the same spectral decomposition procedure was computed based on the memory recall phase of each list, using the 500-ms interval preceding a response vocalization. On matched no-stimulation lists, decoding was performed identically but stimulation was disabled. Full documentation of the closed-loop procedure is described in previous works (6, 7, 14).

<u>Random stimulation:</u> Similar stimulation parameters to those used in the closed-loop were applied to the random stimulation cohort, with the exception of stimulation timing and duration. For 7 patients in the random cohort, stimulation timing and duration were identical to the closed-loop cohort, except that stimulation was triggered using a memory classifier that had been trained on a randomly permuted version of the participant’s record-only data. This meant that classifiers for these participants were at chance in differentiating between good and poor memory encoding states. For the remaining 8 of the patients in the random cohort, stimulation was triggered 200 ms prior to the onset of a word in the block and lasted for 4.6 seconds, until the end of the presentation of the adjacent word in the list. Hence, stimulation occurred in alternating two-word blocks within each list(7, 14).

**Analysis of memory performance:** Recall performance change following stimulation was calculated by previous study(14) , as: Δ 100, where is the average recall for stimulation lists, and is the average recall for no-stimulation lists. Because the first three lists of every stimulation session were always non-stimulated (used for normalization of the classifier input features for that session), these lists were excluded from the calculation of to avoid introducing a temporal order confound(6). All participants were required to demonstrate a minimum = 8.33% (1 out of 12 words per list) for inclusion in the sample.

### Stimulation location and WMP

The coordinates of the radiodense electrode contacts were first obtained from the post-implantation CT and rigidly co-registered to the corresponding volumetric T1-weighted MRI. In the second step, the fused CT-MRI image was non-linearly normalized to MNI space using ANTs(49). Two neuro-radiologists reviewed cross-sectional images and surface renderings.

Using FreeSurfer-based segmentation of each participant’s T1-weighted MRI, we reconstructed the cortical gray–white matter boundary surface. Each stimulation site was defined as the midpoint of the bipolar electrode pair. WMP was quantified as the shortest distance from the stimulation site to the gray–white matter interface, computed by projecting the site onto the local surface normal of the reconstructed boundary in native T1 space, following previously established methods(14).

In MNI space, the stimulation site was modeled as a sphere centered at the midpoint of the bipolar electrode pair with a 12 mm radius, consistent with estimates by Collavini et al. (50) showing that typical clinical currents (∼1–3 mA) produce comparable spatial extents in realistic head models and previous practice(42, 43).

### Localizing stimulation to brain networks

When memory changes are elicited by stimulation at different deep electrode sites across patients, it can be difficult to attribute the effect to a specific brain region, especially when these sites are anatomically distinct. To address this, stimulation sites were integrated with network-level maps derived from diffusion-based structural connectivity and BOLD-based functional connectivity. This network-based approach, originally developed as a “lesion network mapping” technique to interpret the effects of differently located lesions(41), is utilized here to determine the cortical circuits engaged by distinct brain stimulation sites(40, 43). By inserting stimulation site as the “lesioned” area, we enabled the reliable identification of distinct regions, each sourced to a known stimulation site, that form structural and functional networks.

Structural connectivity from the stimulation sites was estimated using the Network Modification (NeMo) Tool v2 (51). This tool leverages a normative tractography database derived from whole-brain probabilistic tractography performed on 420 unrelated healthy subjects from the Human Connectome Project. Streamlines were generated using the iFOD2 algorithm with Anatomically Constrained Tractography (ACT), providing a high-resolution estimate of probabilistic fiber pathways. Each stimulation region, defined in MNI space, was input to the NeMo model as a “lesion mask,” allowing estimation of its impact on whole-brain structural connectivity across a 438-region atlas comprised of 358 cortical parcels from the HCP multimodal parcellation, along with subcortical and cerebellar regions. This mask was non-linearly registered to each subject’s diffusion space to identify streamlines intersecting the region. For each subject, pairwise ChaCo (Change in Connectivity) ratios were calculated, reflecting the proportion of streamlines between each pair of parcels that intersect the stimulation region.

These values were then averaged across all subjects to generate a normative profile of structural connection. Finally, for each parcel, we computed a cumulative ChaCo score by summing its pairwise ChaCo ratios across all other regions, quantifying the relative degree of structural connection between the stimulation region and each brain area in the 438-region atlas.

Functional connectivity was estimated using a seed-based approach, based on a normative dataset (n = 34) collected at Thomas Jefferson University during a word list verbal encoding task. During scanning, HPs were visually presented with 100 single words, grouped into 10 blocks.

Each word was displayed for 3 seconds with an inter-word interval of 1.65 seconds, and an inter-block interval of 15 seconds. To assure compliance and task engagement, participants responded to each word using a keypad to indicate whether they judged the word to be pleasant or unpleasant. Functional MRI data were acquired by a 10-minute session using a Siemens Prisma 3T scanner with a multiband EPI sequence (TR = 0.8 s). fMRI data were preprocessed using *fMRIPrep* 22.1.1(52), which included motion correction, spatial normalization to MNI space, and anatomical segmentation. Postprocessing was performed using XCP-D(53), applying the standard 36-parameter confound regression pipeline and temporal filtering between 0.01–0.08 Hz. We extracted the fMRI task time series data from each stimulated region (in MNI space), as well as from each parcel of atlas (438 total regions) in each healthy participant. Functional connectivity was estimated using Pearson correlations between the stimulation region and each atlas parcel during the verbal encoding task. For each stimulation region, the resulting connectivity maps were averaged across all HPs to derive its normative functional connectivity profile. Notably, task design regressors were intentionally not removed to preserve task-evoked coactivation effects and examine connectivity under naturalistic task engagement.

### Statistics

All statistical comparisons or correlations were conducted as two-tailed tests. Correlations were used to examine the relationship between stimulation-related memory change and the structural or functional connectivity metrics (i.e., the values quantifying the connectivity between the stimulation region and each brain area). The distribution of structural connectivity values was zero-inflated and markedly non-normal, Kendall τ-b correlations were employed. This non-parametric coefficient quantified the strength of monotonic associations while providing robust handling of zero values (no connectivity) and skewed distributions (54–56). Functional connectivity values, which approximated a normal distribution, were analyzed using Pearson correlation.

To control the family-wise error rate across the 438 brain regions, we used a max-statistic (tmax/joint) permutation procedure implemented in PERMUTOOLS (5,000 permutations)(57). On each permutation, stimulation-related memory-change scores were randomly reassigned across participants (connectivity values held fixed), the connectivity–memory association (Pearson’s r) was recomputed for all 438 regions, and the maximum absolute test statistic across regions was retained to form a null distribution of maximal statistics. Permutation-corrected p-values were obtained by comparing the observed regional statistics against this max-statistic null distribution, and regions with corrected p < 0.05 were considered significant. (58–60).

### Stimulation–verbal encoding memory network congruence predicts memory response

We determined how the structural connectivity profile of each stimulation site aligned with a normative verbal-encoding network and, subsequently, whether this alignment predicted stimulation-related memory performance. To accomplish this, the normative verbal-encoding network was derived from fMRI memory task data collected from 34 HPs (see above for a description of this verbal memory encoding task). After preprocessing with fMRIPrep (see Functional Connectivity above), first-level GLM analyses were performed in SPM12 using an 8 mm FWHM smoothing kernel and a 24-parameter motion regression model. A *word > rest* contrast was computed for each participant, and the resulting t-statistic maps were averaged to obtain a group-level verbal encoding activation pattern. This activation map was then parcellated into the same 438-region atlas used for our structural connectivity analyses, yielding a parcelwise normative verbal-encoding network that could be directly compared to stimulation-related structural connectivity profiles.

To illustrate the analytical framework, we selected a representative pair of thresholds. First, the verbal-encoding network map was the top 30% of positively weighted parcels. Second, each patient’s structural connectivity map from the stimulation site was thresholded at 1, which corresponds to approximately the 80^th^-90^th^ percentile of that patient’s structural connectivity distribution. For each thresholded pair of maps, we binarized parcels and quantified their spatial overlap using the Dice similarity coefficient. Dice is a normalized measure based on the size of the intersection relative to the combined extent of the two binary masks. In this context, it can be interpreted as the degree to which a stimulation site is structurally coupled to the same regions that showed strong task-related verbal-encoding activation at the group level. Therefore, this representative threshold pair highlighted the regions that are both strongly engaged during verbal encoding and strongly anatomically connected to the stimulation site, providing an intuitive visualization of network congruence. We then computed Spearman correlations between Dice coefficients and stimulation-related memory changes, separately for the closed-loop and random-stimulation cohorts.

We then evaluated the robustness of this relationship across a broader parameter space. We systematically varied the verbal-encoding network threshold from 0.1 to 1.0 (ten levels) and the structural connectivity threshold from 0.001 to 50 (six levels) to construct a two-dimensional parameter grid. For each pair of thresholds, we repeated the above procedure. Multiple comparisons across the full grid were controlled using the Benjamini–Hochberg false discovery rate (FDR; q < 0.05).

### Partial least squares structural equation modeling

Partial least squares structural equation modeling (PLS-SEM; SmartPLS 4) was performed to examine the multivariate relationships among connectivity-derived latent variables, stimulation mode, baseline memory (word lists presented with no stimulation), white matter proximity, and stimulation-related memory changes. This variance-based SEM approach is well-suited for exploratory neuroimaging data because it does not assume multivariate normality and accommodates complex mediation or moderation structures in modest sample sizes. As the selected structural connectivity regions contained a small proportion of zero values, we replaced zeros with the minimal non-zero value and applied a natural logarithmic transformation to approximate a normal distribution prior to model inclusion. Model estimation was conducted using a path-weighting scheme with 5,000 bootstrap resamples to obtain significance estimates for the outer loadings and the direct and indirect path coefficients.

All models were evaluated according to standard PLS-SEM quality criteria, including indicator reliability, internal consistency, average variance extracted (AVE > 0.50), and discriminant validity (heterotrait–monotrait ratio < 0.85). Path coefficients were interpreted based on bias-corrected bootstrap confidence intervals and considered significant at *p* < 0.05.

### Data and Code Availability

Anonymized data, analysis code, statistical results, and brain maps generated in this study are publicly available at https://github.com/HerzLab/ConnectivityInMemoryStimulation. Structural connectivity measures were computed using the Network Modification (NeMo) Tool v2 (https://github.com/kjamison/nemo). Functional MRI data processing was performed using *fMRIPrep* 22.1.1(https://fmriprep.org/en/stable/) and XCP-D(https://xcp-d.readthedocs.io). Permutation-based statistical testing was performed with PERMUTOOLS (https://github.com/mickcrosse/PERMUTOOLS). Partial least squares structural equation modeling was conducted using SmartPLS 4 (https://www.smartpls.com). Brain surface visualizations were generated using the ENIGMA Toolbox (https://enigma-toolbox.readthedocs.io/).

## Acknowledgments

This work was supported by the National Institute of Neurological Disorders and Stroke of the U.S. National Institutes of Health (R01 NS112816-01, J.I.T.), the American Epilepsy Society Postdoctoral Research Fellowship (1277380, Q.Z.).

## Competing Interests

The authors declare no competing interests.

## Notes

https://github.com/HerzLab/ConnectivityInMemoryStimulation

### Competing Interest Statement

The authors have declared no competing interest.

