## Supplementary Fig. S1 for "Structural and Functional Connectivity Predict the Effects of Direct Brain Stimulation on Memory"

Supplementary table 1. Sample demographic characteristics

| **Subject**  **ID** | **Stimulation type** | **Sex** | **Age** | **Subject ID** | **Stimulation type** | **Sex** | **Age** |
| --- | --- | --- | --- | --- | --- | --- | --- |
| R1154D | closed-loop | Female | 36.14 | R1378T | closed-loop | Male | 38.80 |
| R1195E | closed-loop | Male | 44.37 | R1379E | closed-loop | Male | 49.75 |
| R1200T | closed-loop | Male | 25.97 | R1380D | closed-loop | Female | 49.21 |
| R1204T | closed-loop | Female | 25.77 | R1383J | closed-loop | Male | 25.92 |
| R1217T | closed-loop | Male | 37.25 | R1385E | closed-loop | Female | 33.84 |
| R1226D | closed-loop | Female | 41.75 | R1390M | closed-loop | Male | 32.07 |
| R1230J | closed-loop | Female | 56.26 | R1391T | closed-loop | Male | 29.04 |
| R1236J | closed-loop | Female | 51.73 | R1398J | closed-loop | Male | 45.30 |
| R1243T | closed-loop | Male | 63.92 | R1406M | closed-loop | Female | 52.20 |
| R1260D | closed-loop | Female | 57.00 | R1465D | closed-loop | Male | 35.10 |
| R1264P | closed-loop | Female | 52.59 | R1477J | closed-loop | Male | 19.13 |
| R1274T | closed-loop | Female | 44.34 | R1487T | closed-loop | Male | 54.16 |
| R1286J | closed-loop | Female | 57.58 | R1488T | closed-loop | Male | 50.75 |
| R1292E | random | Female | 39.32 | R1489E | closed-loop | Female | 32.46 |
| R1308T | random | Female | 40.08 | R1498D | closed-loop | Male | 30.03 |
| R1315T | random | Male | 22.38 | R1050M | random | Male | 20.41 |
| R1317D | random | Male | 36.50 | R1111M | random | Male | 20.06 |
| R1323T | random | Male | 40.17 | R1163T | closed-loop | Male | 45.68 |
| R1330D | random | Female | 37.80 | R1170J | closed-loop | Male | 20.62 |
| R1334T | random | Female | 38.14 | R1176M | random | Female | 41.38 |
| R1339D | random | Male | 29.24 | R1177M | random | Female | 23.39 |
| R1341T | random | Male | 32.71 | R1180C | random | Female | 21.45 |
| R1345D | random | Male | 48.22 | R1223E | closed-loop | Female | 42.75 |
| R1374T | closed-loop | Male | 60.16 | R1235E | closed-loop | Male | 48.73 |
| R1375C | closed-loop | Female | 22.97 | R1293P | closed-loop | Male | 38.09 |

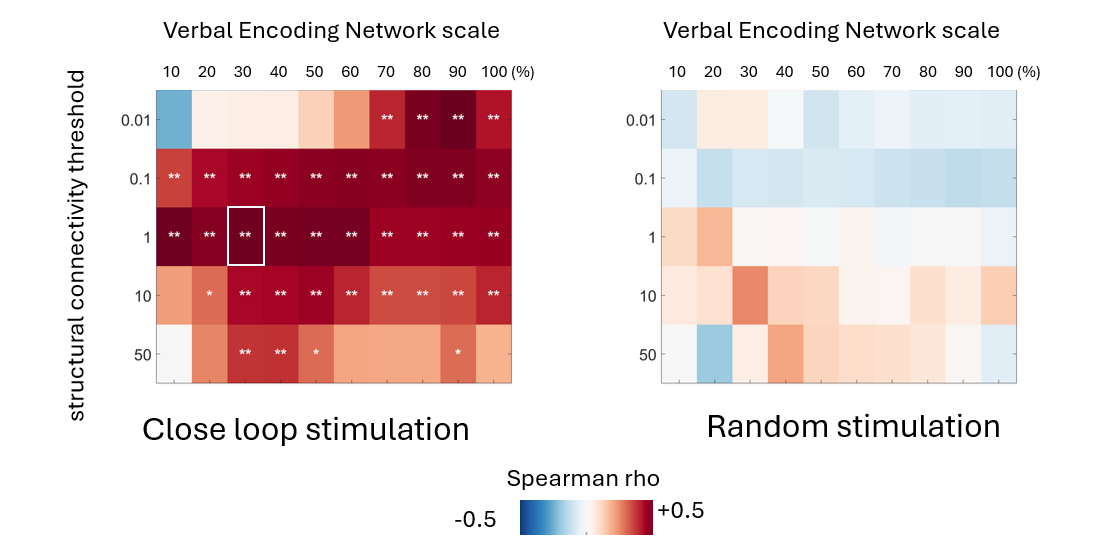

**Figure S1. Structural connectivity–memory associations are robust across threshold parameters.** We systematically varied both the verbal encoding network (retaining the top 10%–100% of voxels within the normative verbal encoding network) and structural connectivity (0.001–50, reflecting the magnitude of stimulation-related structural connectivity strength) thresholds to generate a two-dimensional parameter grid for each stimulation condition. For each threshold pair, we computed the correlation between network congruence (Dice coefficient) and stimulation-related memory change. Left panel: The closed-loop condition showed a consistent positive association across a broad range of thresholds. Figure 3d used a representative operating threshold corresponding to the maximum correlation values in the matrix (the white-boxed cell). Right: No significant structural-memory associations were observed for the random stimulation condition. Multiple comparisons were controlled using FDR (q < 0.05). * q < 0.05; ** q < 0.01.
